# Cell-Type-Resolved Pseudobulk Classification Across Independent Cohorts Identifies Microglial PTPRG as a Transcriptional Hub in Alzheimer’s Disease

**DOI:** 10.64898/2026.04.07.717029

**Authors:** Agata Marchi, Danish Anwer, Eduard Kerkhoven, Nicola Pietro Montaldo, Jeroen Gilis, Annikka Polster

## Abstract

Alzheimer’s disease (AD) is a progressive neurodegenerative disorder characterized by cognitive decline and widespread cerebral pathology. Understanding cell-type-specific molecular mechanisms underlying AD is critical for identifying precise therapeutic targets.

We applied a supervised machine learning approach to single-nucleus RNA sequencing data from the ROSMAP cohort, aggregating gene expression profiles into pseudobulk representations across six major brain cell types.

Systematic evaluation of all possible cell-type combinations identified microglia and astrocytes as the most discriminative cell types for AD classification. A logistic regression model trained on 228 highly variable genes achieved robust classification performance on held-out ROSMAP samples (balanced accuracy 0.87, AUC 0.89) and generalized to an independent cohort from the Seattle Alzheimer’s Disease Brain Cell Atlas (balanced accuracy 0.86, AUC 0.92), demonstrating cross-cohort reproducibility that remains uncommon in computational AD research. Among the 72 genes selected by the model, microglial PTPRG exhibited the highest absolute coefficient. Gene Set Enrichment Analysis (GSEA) revealed that microglia-expressed genes were enriched for chronic immune activation and inflammatory signaling, while astrocyte-associated genes implicated protein homeostasis stress and HSF1-mediated chaperone pathways. Weighted Gene Co-expression Network Analysis (WGCNA) further showed that PTPRG operates within fundamentally different gene network contexts in AD and NCI microglia, with AD networks characterized by inflammatory dysregulation and NCI networks reflecting homeostatic immune surveillance. Cell-cell communication analysis identified established AD risk genes including APOE, GRN, PSEN1, and CLU among the top neuronal ligands predicted to regulate microglial PTPRG, positioning it as a convergence point for disease-relevant neuronal signals. Correlation analysis further revealed that excitatory and inhibitory neurons couple to microglial PTPRG through distinct biological processes, implicating divergent mechanisms of AD-associated microglial dysregulation.

Collectively, these findings establish microglial PTPRG as a central hub integrating neuronal signaling and inflammatory dysregulation in AD pathology.

## INTRODUCTION

Alzheimer’s disease (AD) is the most common cause of dementia, characterized by progressive cognitive decline and widespread neurodegeneration (DeTure and Dickson 2019). Despite extensive research, the molecular mechanisms underlying disease onset and progression remain incompletely understood, and the cellular complexity of the human brain presents a fundamental challenge: AD affects multiple neuronal and non-neuronal cell types, each contributing differently to disease pathophysiology (Mathys et al., 2019; Saura et al., 2023; Smith et al., 2022; Luquez et al., 2022). Resolving cell-type-specific molecular alterations in human brain tissue is therefore essential for advancing mechanistic understanding and identifying precise therapeutic targets.

Early bulk RNA sequencing studies provided valuable insights into region-specific gene expression patterns in AD and enabled molecular subtyping and target identification (Twine et al., 2011; Neff et al., 2021). However, by averaging expression across heterogeneous cell populations, bulk approaches obscure cell-type-specific contributions to disease pathophysiology. Single-nucleus RNA sequencing (snRNA-seq) addresses this limitation directly, enabling systematic characterization of transcriptional changes across individual brain cell populations and revealing distinct disease-associated microglial and astrocytic subpopulations that emerge in response to AD pathology (Habib et al., 2020; Olah et al., 2020; Deczkowska et al., 2018; Mathys et al., 2024). This resolution is particularly important given the selective vulnerability of specific neuronal subpopulations to tau pathology, including RORB-expressing excitatory neurons in the entorhinal cortex that are depleted early in disease progression, and the dynamic, cell-type-specific transcriptional changes that accompany AD across the full spectrum of brain cell classes (Leng et al., 2021; Luquez et al., 2022).

Yet the same cellular heterogeneity that motivates snRNA-seq also complicates the identification of disease-associated transcriptional signatures. AD pathology does not affect all cells or brain regions equally (Fu et al., 2018), meaning that not all cells in a diagnosed individual necessarily exhibit molecular changes linked to disease. Differential expression approaches can characterize these changes descriptively, but supervised learning is needed to systematically identify which molecular features are most discriminative between disease states and to quantify their relative contributions across cell types.

Applying supervised learning to single-cell data, however, introduces a labeling problem: clinical diagnoses operate at the sample level, and extending these labels to individual cells is methodologically unsound, since we cannot determine which specific cells are affected or to what degree. Several approaches have been proposed to address this. Craig et al. (2025) identify disease-associated cells from sample-level labels alone, but treat all cells from healthy donors as a uniform baseline. He et al. (2024) use an attention-based model that avoids cell-level label assignment, but analyzes each cell type independently and therefore cannot capture disease mechanisms arising from coordinated changes across cell types. Verlaan et al. (2025) create unified sample-level representations by pooling across heterogeneous cell populations, successfully avoiding incorrect labeling, but at the cost of discarding cell-type identity entirely, and their approach performs on par with methods that aggregate all cells without any cell-type separation.

We instead adopt pseudobulk aggregation performed separately for each cell type, representing each sample as a vector of cell-type-specific expression profiles. This preserves cell-type information while avoiding incorrect cell-level labeling, and enables systematic, unbiased comparison of the discriminative contribution of each cell type to AD classification.

Applying this approach to snRNA-seq data from the ROSMAP cohort and independently validating findings in the Seattle Alzheimer’s Disease Brain Cell Atlas (SEAD), we identify microglia and astrocytes as the cell types carrying the strongest transcriptional signal of AD, and establish microglial PTPRG as a central hub integrating neuronal signaling and inflammatory dysregulation in disease pathology.

## METHODS

### Data pre-processing

Six major cell types were included in the analysis: microglia, astrocytes, excitatory neurons, inhibitory neurons, oligodendrocyte precursor cells, and oligodendrocytes. Preprocessing was performed independently for each.

At the cellular level, quality control filtering was performed using the *scanpy* package (Wolf et al., 2018). Cells expressing at least 200 genes were retained, while genes were required to be detected in a minimum of 200 cells to be kept. Cells with >5% mitochondrial gene content were excluded to remove low-quality cells. Only individuals with at least 30 cells per cell type were retained for downstream analysis.

At the subject-level we identified and removed 11 outliers based on inconsistent cell type annotations. These outliers were detected by comparing the cell type annotations performed by Mathys et al., (2023) with those we generated using *CellTypist* (Domínguez Conde et al., 2022). Subjects were flagged as outliers when more than 50% of their cells received discordant annotations between the two methods, indicating unreliable cell type assignments that could compromise downstream analyses. After preprocessing and quality control, the ROSMAP cohort comprised 367 subjects.

Subject phenotypic labels were reassigned based on the combination of clinical consensus diagnosis of cognitive status at time of death (*cogdx*) (Schneider et al., 2013), CERAD score (*ceradsc,* semiquantitative measure of neuritic plaques), and Braak (Bennett, D A et al., 2006) stage (*braaksc,* semiquantitative measure of neurofibrillary tangles). To identify the most extreme cases, the same approach as Verlaan et al. (2025) was followed in order to ensure that the gene expression patterns of AD samples would be as distinct as possible from those of NCI samples.

The classification criteria were as follows:

● Subjects meeting stringent pathological and cognitive decline criteria (*cogdx = 4, braaksc>=4, ceradsc <=2*) were classified as Alzheimer’s Disease (AD).
● Subjects with no cognitive impairment and minimal pathology (*cogdx = 1, braaksc<=3, ceradsc>=3*) were classified as Not Cognitively Impaired (NCI).
● Subjects with mild cognitive impairment and neuritic plaques (cogdx = 2, braaksc >= 4, ceradsc <= 2) were classified as Mildly Cognitively Impaired with Plaques (MCI+P)
● All remaining subjects were classified as “unknown” and excluded from further analysis.

Following phenotypic label reassignment based on the composite classification scheme, the cohort was partitioned into two datasets: the MCI+P dataset consisting of 32 subjects and the AD-NCI dataset consisting of 148 subjects (84 AD, 64 NCI).

### Pseudobulk aggregation and dataset construction

To preserve the cell-type specificity of each gene and enable direct comparison of expression patterns across different cell populations, gene expression values were aggregated to the pseudobulk level for each cell type separately by summing expression values across all cells for each subject, following count depth scaling to 10^6^. Pseudobulk expression matrices from different cell types were concatenated to produce a subject-by-feature matrix where each row represented an individual and columns contained pseudobulk expression values for each gene-cell type combination. This concatenation procedure was performed independently for the AD-NCI dataset and the MCI+P dataset.

### External validation cohort

The Seattle Alzheimer’s Disease Brain Cell Atlas (SEAD) (Gabitto et al., 2024) dataset was used as an independent external test cohort to evaluate model generalizability. The SEAD dataset underwent identical preprocessing steps as the ROSMAP cohort. To ensure compatibility between datasets, both ROSMAP and SEAD were filtered to retain only genes present in corresponding cell types across both cohorts. Only subjects meeting the AD or NCI definitions were retained. This resulted in 48 samples, 39 labeled as AD and 9 as NCI.

### Classification of AD and NCI groups

The AD-NCI dataset was split into training (80%) and testing (20%) sets. We then used the training set to identify which cell types, alone or in combination, contribute most strongly to disease classification, as well as the genes that provide the most informative features.

To evaluate the contribution of different cell types, we analyzed every possible combination of the six cell types, from single cell types to all six combined (63 total combinations).

For each cell-type combination, we trained a logistic regression classifier using 5-fold cross-validation. Within each cross-validation iteration, highly variable genes (HVGs) were selected in the train fold using the *highly_variable_genes* function in scanpy. HVGs were selected separately for each cell type included in the combination. Different thresholds for HVGs selection were evaluated: 100, 300, 500, 700, 1000.

The model performance was evaluated on each fold using balanced accuracy, F1 score, and area under the ROC curve (AUC).

To identify the optimal combination of cell types, we considered both the mean balanced accuracy achieved by each combination across folds and the number of selected features, aiming to balance predictive performance with model complexity. To ensure robust and reproducible feature selection, we retained only genes that were consistently identified as highly variable across all five cross-validation folds for the chosen cell-type combination. The intersection was defined as the final feature set and used to train a final classification model on the full training dataset.

The final L1-regularized logistic regression model was trained on the complete training set using the selected features, the model’s performance was then evaluated on the ROSMAP held-out test set and on the independent SEAD cohort (Lein et al., 2024).

The contribution of individual genes to disease classification was determined by the absolute values of non-zero coefficients from the final L1-regularized logistic regression model. Gene Set Enrichment Analysis (GSEA) was subsequently performed on this gene set using the Enrichr framework implemented in GSEApy (Fang et al., 2023), querying the MSigDB Hallmark 2020, KEGG 2021 Human, and Reactome Pathways 2024 databases.

The final trained model was also used to estimate the position of the MCI+P samples along the AD-NCI spectrum by computing the predicted probabilities of AD.

### Weighted Gene Co-expression Network Analysis

Raw single-nucleus RNA sequencing data from microglial cells were preprocessed as described above and aggregated into pseudobulk profiles by summing raw counts across all cells for each subject. Counts were normalized to counts per million and log1p-transformed to stabilize variance prior to Weighted Gene Co-expression Network Analysis (WGCNA), performed independently for AD and NCI groups using PyWGCNA (Rezaie et al., 2023).

Co-expression modules were identified via dynamic tree cutting and merged based on eigengene similarity. GSEA was subsequently performed on the genes within the PTPRG-containing module separately for each condition, using the Enrichr framework in GSEApy (Fang et al., 2023) and querying the MSigDB Hallmark 2020, KEGG 2021 Human, and Reactome Pathways 2024 databases.

### Cell-cell communication using NicheNet

To determine whether the observed differences in the gene network surrounding PTPRG translate into altered signaling between neurons and microglia, we performed cell-cell communication analysis using NicheNet (Browaeys et al., 2020). This framework integrates prior knowledge of signaling and gene regulatory networks to predict how ligands expressed by sender cells influence gene expression in receiver cells.

The analysis was performed separately for excitatory and inhibitory neurons. Differentially expressed genes in microglia were first identified by comparing AD and NCI samples using the NicheNet pipeline with a case-control design (adjusted p ≤ 0.001, |log fold-change| ≥ 0.25). Ligand-receptor interaction networks were then constructed using NicheNet’s prior knowledge databases, with expressed genes defined by a 5% expression threshold in both sender (neurons) and receiver (microglia) cells. Ligand activity analysis was performed to identify neuronal ligands best explaining the microglial DEGs, with ligands ranked by their area under the precision-recall curve. Finally, ligand-receptor-target interactions were identified with PTPRG as the target, and corresponding regulatory potential scores were calculated.

GSEA was subsequently performed on the sets of ligands predicted to regulate microglial PTPRG, separately for excitatory and inhibitory neuron-microglia pairs, using all identified ligands as background. The analysis was conducted via the Enrichr framework in GSEApy (Fang et al., 2023), querying the MSigDB Hallmark 2020, KEGG 2021 Human, and Reactome Pathways 2024 databases.

### Gene expression correlation analysis between microglial and neuronal cells

Linear regression models were fitted separately for excitatory and inhibitory neurons to evaluate the association between neuronal gene expression and microglial PTPRG expression while controlling for potential confounders. Each model was specified as:

> *neuronal gene expression ∼ PTPRG expression + clinical diagnosis + age at death + sex + post-mortem interval + batch*

where neuronal gene expression and PTPRG expression were log1p-transformed pseudobulk values, clinical diagnosis (AD vs. NCI) and sex were encoded as categorical variables, and age at death and post-mortem interval were included as continuous covariates. Models were estimated using ordinary least squares with robust standard errors. The regression coefficient (β) for PTPRG was extracted for each neuronal gene alongside its p-value, and multiple testing correction was applied using the Benjamini-Hochberg FDR method (adjusted p < 0.05). Only subjects with pseudobulk measurements available for both microglia and the target neuronal population were retained, yielding 119 subjects for the excitatory and 108 for the inhibitory neuron analysis. Genes significantly associated with microglial PTPRG were subsequently subjected to pathway enrichment analysis using GSEApy (Fang et al., 2023) using the Enrichr framework and querying the MSigDB Hallmark 2020, KEGG 2021 Human, and Reactome 2022 databases.

## Results

To identify cell type-specific molecular signatures distinguishing AD from NCI individuals, we applied a logistic regression model to pseudobulk single-nucleus RNA sequencing data from the ROSMAP cohort. The model’s performance was assessed on a held-out ROSMAP test set and validated on an independent SEAD cohort. Subsequently, we conducted feature importance analysis to identify the key genes used for the classification, applied weighted gene co-expression network analysis (WGCNA) to identify and characterize disease-associated gene modules in microglia performed cell-cell communication analysis using NicheNet to infer ligand-receptor interactions, and examined neuron-microglia gene expression correlations to assess the strength of transcriptomic coordination between these cell types in the context of AD pathology.

### Identification of optimal cell-type combination for AD classification

Among the 63 combinations tested, 10 achieved a mean balanced accuracy between 79% and 80% (Supplementary Table S1). We chose to prioritize a simpler model over a 1% gain in accuracy. Accordingly we selected the combination of astrocytes and microglia, with 500 HVGs selected from each cell type. This model achieved a mean balanced accuracy of 79.75% and a mean F1 score of 0.8375 across five-fold cross-validation.

Using this optimal cell type combination, we identified genes that were consistently selected as highly variable across all five cross-validation folds. Of the 500 highly variable genes (HVGs) selected in each individual fold, 228 genes appeared in all five folds, constituting a robust feature set for final model training. We trained an L1-regularized (Lasso) logistic regression model using these 228 genes on the complete ROSMAP training dataset and evaluated performance on the held-out ROSMAP test set. The model achieved strong discriminative performance with a balanced accuracy of 0.87, AUC of 0.89, and F1 score of 0.88 (Figure 1A). The confusion matrix showed 93% sensitivity for AD samples with a false negative rate of only 6.7% (Figure 1B).

**Figure 1.**
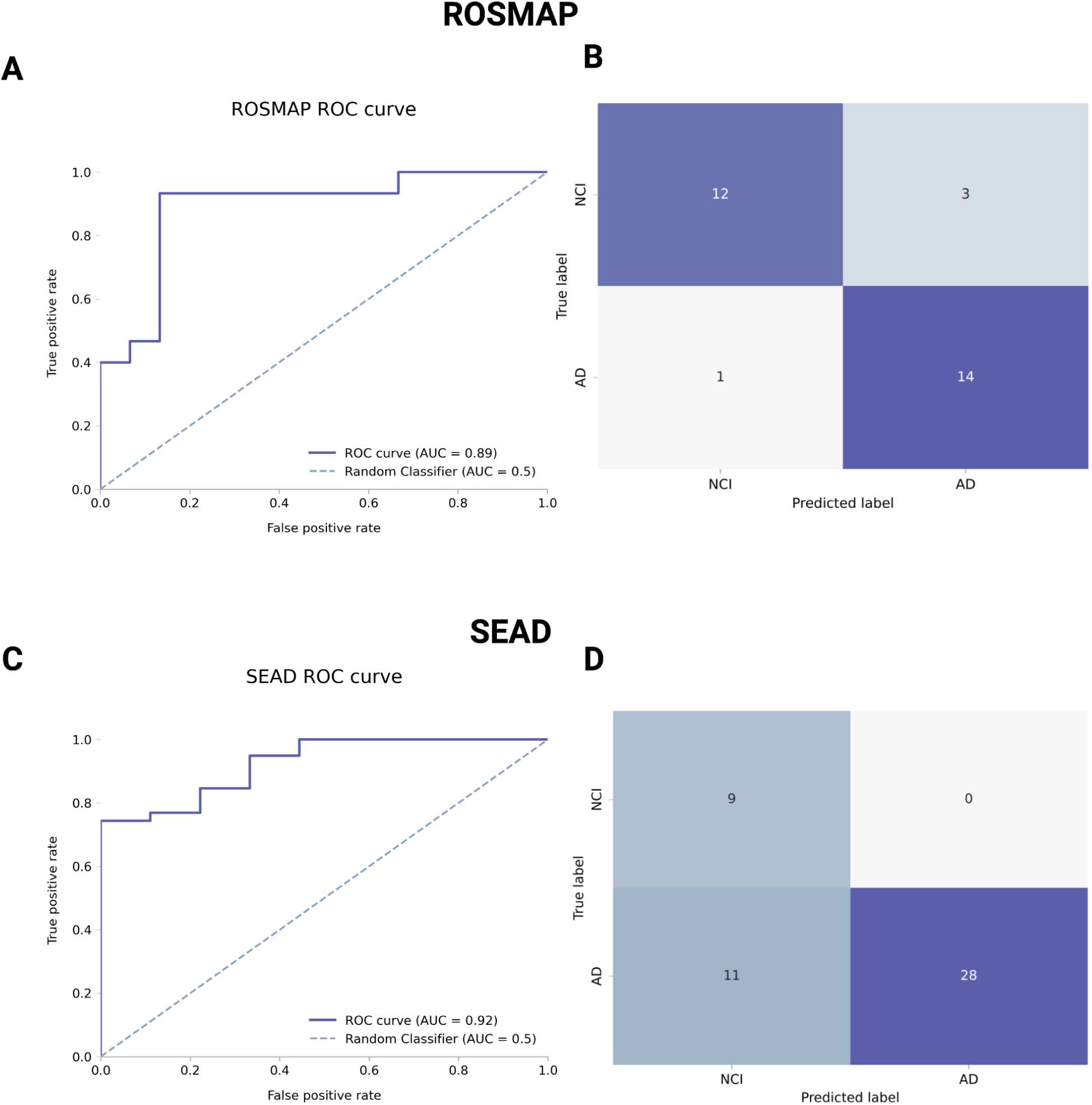
Logistic regression model performance on the ROSMAP test set. and the SEAD cohort. **A)** ROSMAP: Receiver operating characteristic (ROC) curve showing the trade-off between true positive and false positive rates across classification thresholds (AUC = 0.89). **B)** ROSMAP: Confusion matrix displaying the number of correct and incorrect predictions for each class (NCI vs. AD). **C)** SEAD: Receiver operating characteristic (ROC) curve showing the trade-off between true positive and false positive rates across classification thresholds (AUC = 0.92). **D)** SEAD: Confusion matrix displaying the number of correct and incorrect predictions for each class (NCI vs. AD).

To assess generalizability beyond the ROSMAP cohort, we evaluated the model on the independent SEAD validation dataset. The model maintained robust performance with an AUC of 0.92 (Figure 1C), balanced accuracy of 0.86, and F1 score of 0.84. The confusion matrix showed perfect specificity, correctly identifying all NCI samples with no false positives, while achieving 72% sensitivity for AD samples (Figure 1D). These results demonstrate that molecular signatures derived from astrocytes and microglia generalize effectively across independent cohorts.

### Intermediate samples reveal disease continuum

In addition to subjects classified as AD and NCI, the ROSMAP cohort also includes individuals exhibiting mild cognitive impairment and with neuritic plaques. To investigate whether our model trained on extreme cases could capture this disease spectrum, we applied it to MCI+P samples to estimate their position along the AD continuum.

The predicted probabilities for these samples fall between those of the extreme groups, consistent with their intermediate status. This indicates that the model can meaningfully place individuals along the disease continuum, rather than simply distinguishing between two discrete classes (Figure 2).

**Figure 2.**
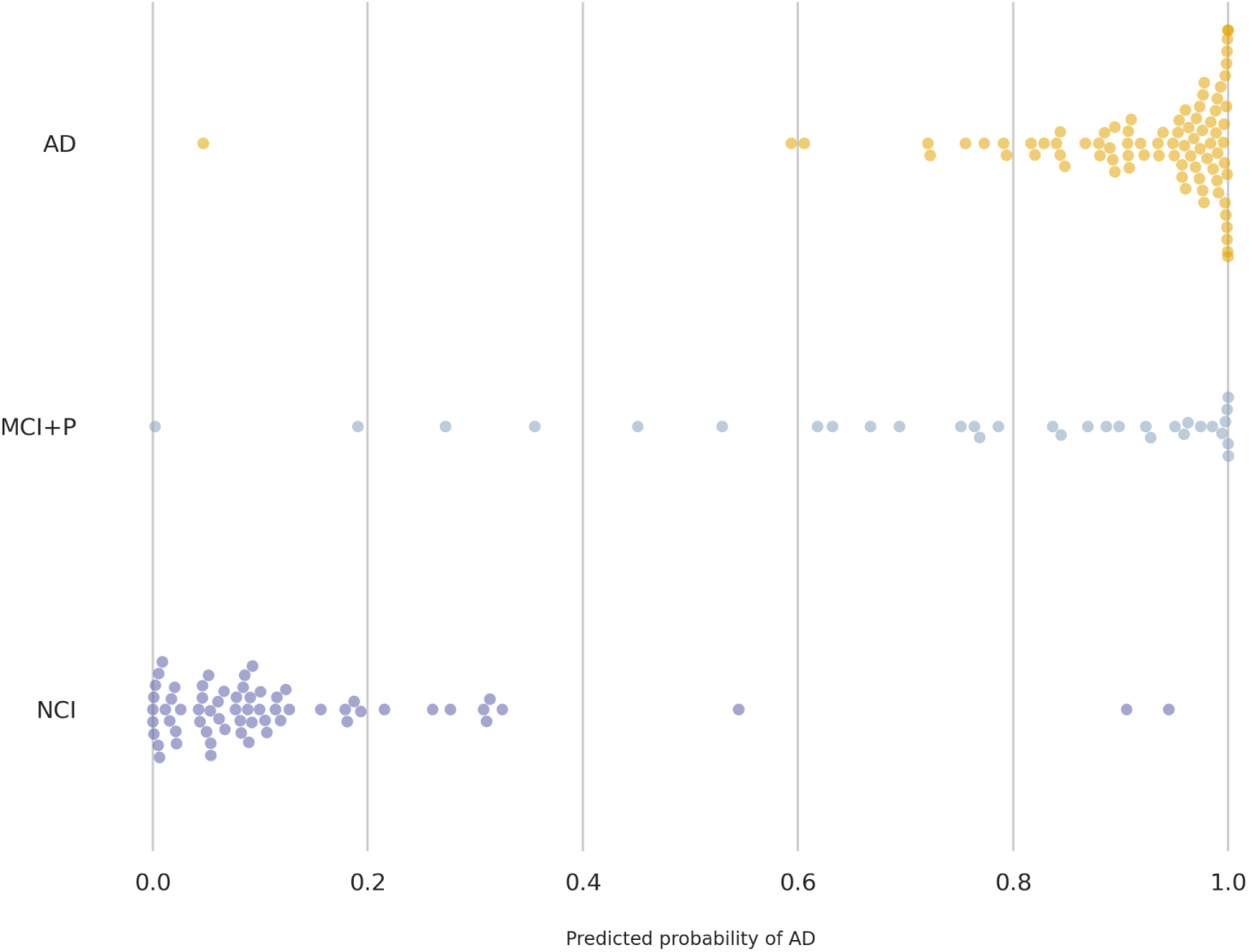
Distribution of predicted AD probabilities across disease severity. Predicted AD probabilities for extreme (AD and NCI) and MCI+P samples from the ROSMAP cohort. Probabilities for MCI+P samples were computed using the logistic regression model trained only on the extreme AD and NCI cases.

### Feature importance analysis reveals key molecular drivers

To identify the genes that contribute most to AD classification, we examined the coefficients of the trained logistic regression model. Among the 228 selected features, L1 regularization resulted in a sparse model with 72 genes having non-zero coefficients. Of these, 47 genes originated from microglia and 25 from astrocytes.

Among the 72 selected important features, PTPRG expressed in microglia was the gene with the highest absolute coefficient value (Figure 3). PTPRG is a protein tyrosine phosphatase gene that acts as a critical tumor suppressor involved in regulating cell growth, differentiation, and preventing malignant transformation (Boni et al., 2022).

**Figure 3.**
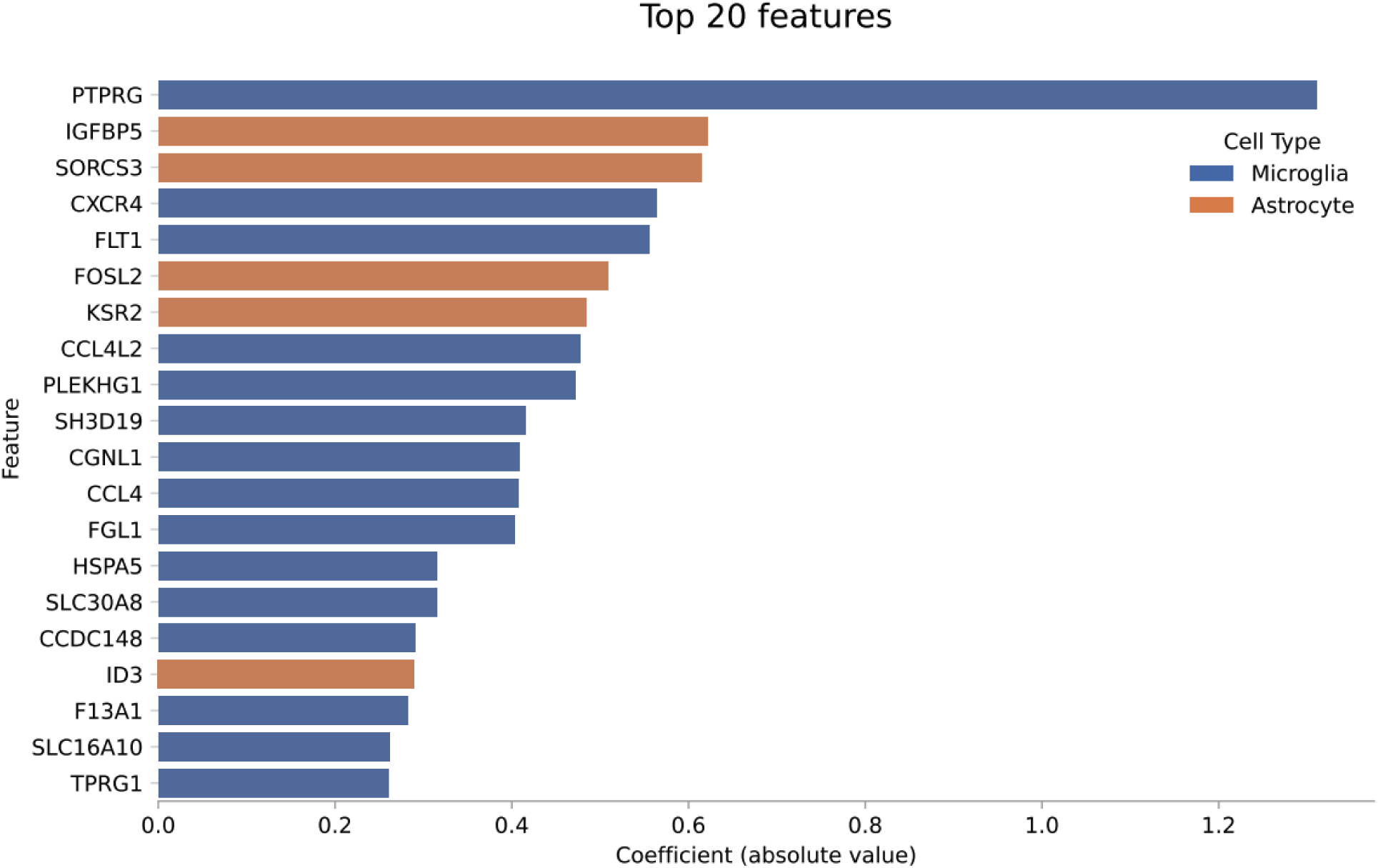
Logistic regression features importance. Top 20 features for the final logistic regression model ranked by absolute coefficient values.

Beyond PTPRG, key microglial features included CXCR4, a chemokine receptor implicated in neuroinflammation, FLT1, which drives chemotactic inflammatory responses, and several additional chemokines and immunoregulatory factors. Among astrocyte-derived features, the top contributors were IGFBP5, a modulator of IGF signaling, and SORCS3, which is involved in cell-surface receptor localization.

While the selected features showed complex expression patterns across samples (Figure 4A), the microglial genes PTPRG and FLT1 exhibited statistically significant different expression levels (Wilcoxon rank-sum test, FDR corrected p-value < 0.05) between the AD and the NCI groups in the test set (Figure 4B and Figure 4C).

**Figure 4.**
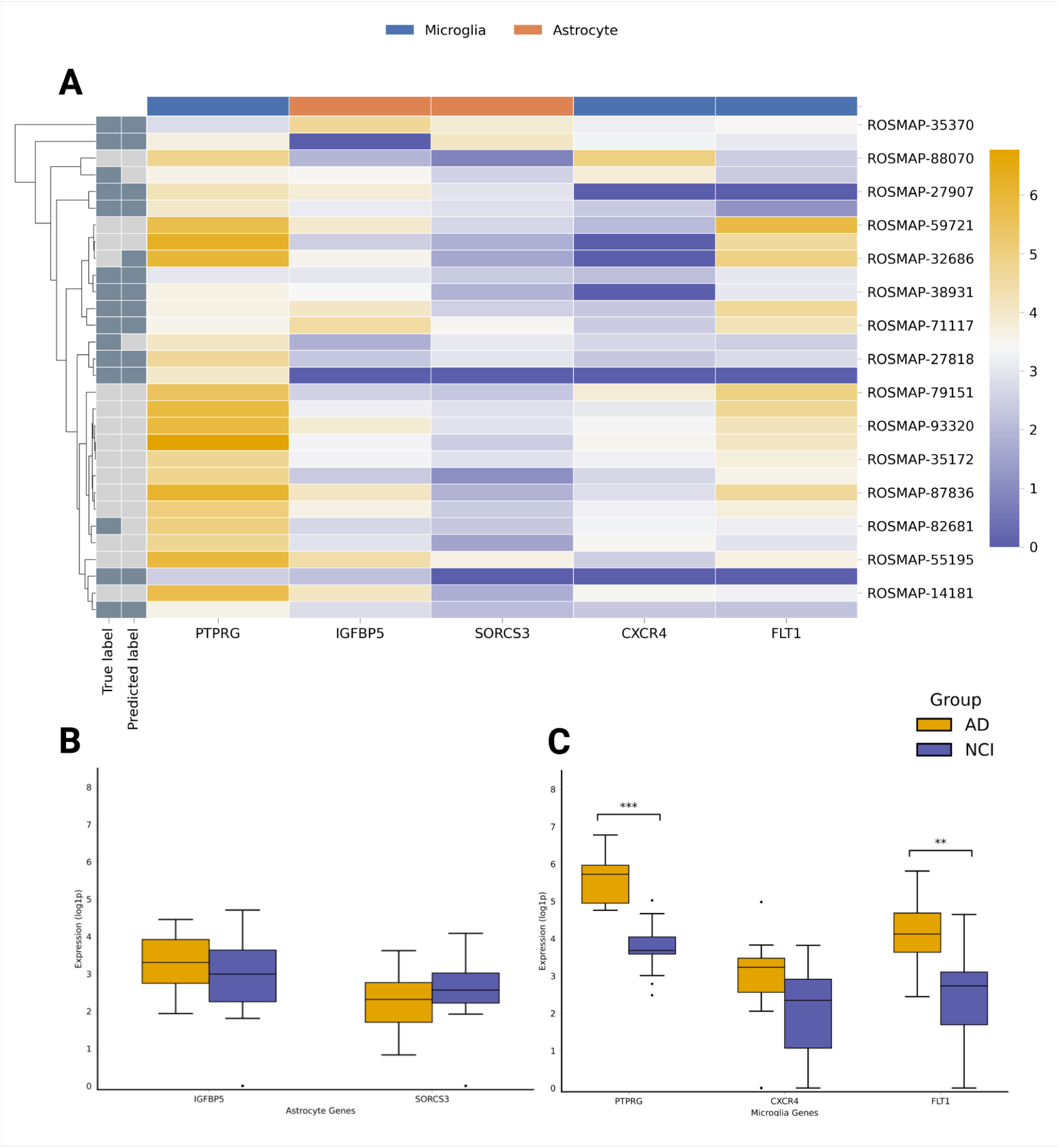
Expression patterns of the top 5 genes with highest feature importance in the logistic regression model. Expression values are shown at the pseudobulk level (library size normalized to 10^6^, log1p transformed). **A)** Heatmap showing expression levels across all test set samples. The leftmost column displays the true diagnostic labels (dark gray: NCI, light gray: AD) alongside the label predicted by the model (predicted label). PTPRG in Microglia shows a clear dichotomous pattern between the two groups. **B)** Boxplots comparing expression distributions between AD and NCI samples for the top genes in Astrocytes. **C)** Boxplots comparing expression distributions between AD and NCI samples for the top genes in Microglia. Statistically significant differences are indicated by asterisks: * p<0.05, ** p<0.01, *** p<0.001.

To investigate the biological processes associated with the 72 most predictive genes, we performed GSEA separately for the microglia-associated and astrocyte-associated gene sets. Microglia (47 genes) exhibited a strongly reactive phenotype characterized by chronic immune activation and ongoing inflammatory signaling alongside compensatory cell-protective stress responses (Figure S1). Astrocytes (25 genes) also showed hallmarks of inflammation as well as protein-homeostasis stress, with enrichment of pathways related to HSF1-mediated responses and chaperone pathways (Figure S2).

Together, the microglia and astrocyte gene sets point to a coordinated glial response in AD, characterized by inflammatory dysregulation in microglia and proteostasis stress in astrocytes.

### Gene Co-expression Networks reveal disease-specific disruption of PTPRG coordination

To investigate whether the gene network surrounding PTPRG is altered in AD, we performed WGCNA separately on microglial cells from AD and NCI samples. The two analyses yielded distinct co-expression module structures: the AD network produced 20 final modules (merged from an initial 47), while the NCI network produced 14 (merged from an initial 38) (Figure 5A). Differences in network architecture were also reflected in the soft-thresholding parameters, with the AD network requiring a lower power than the NCI network (8 vs. 10, both at R² = 0.9), and exhibiting substantially lower mean connectivity (3.64 vs. 8.56). Despite comparable scale-free topology fits, these parameters indicate a more fragmented co-expression structure in AD relative to NCI.

**Figure 5.**
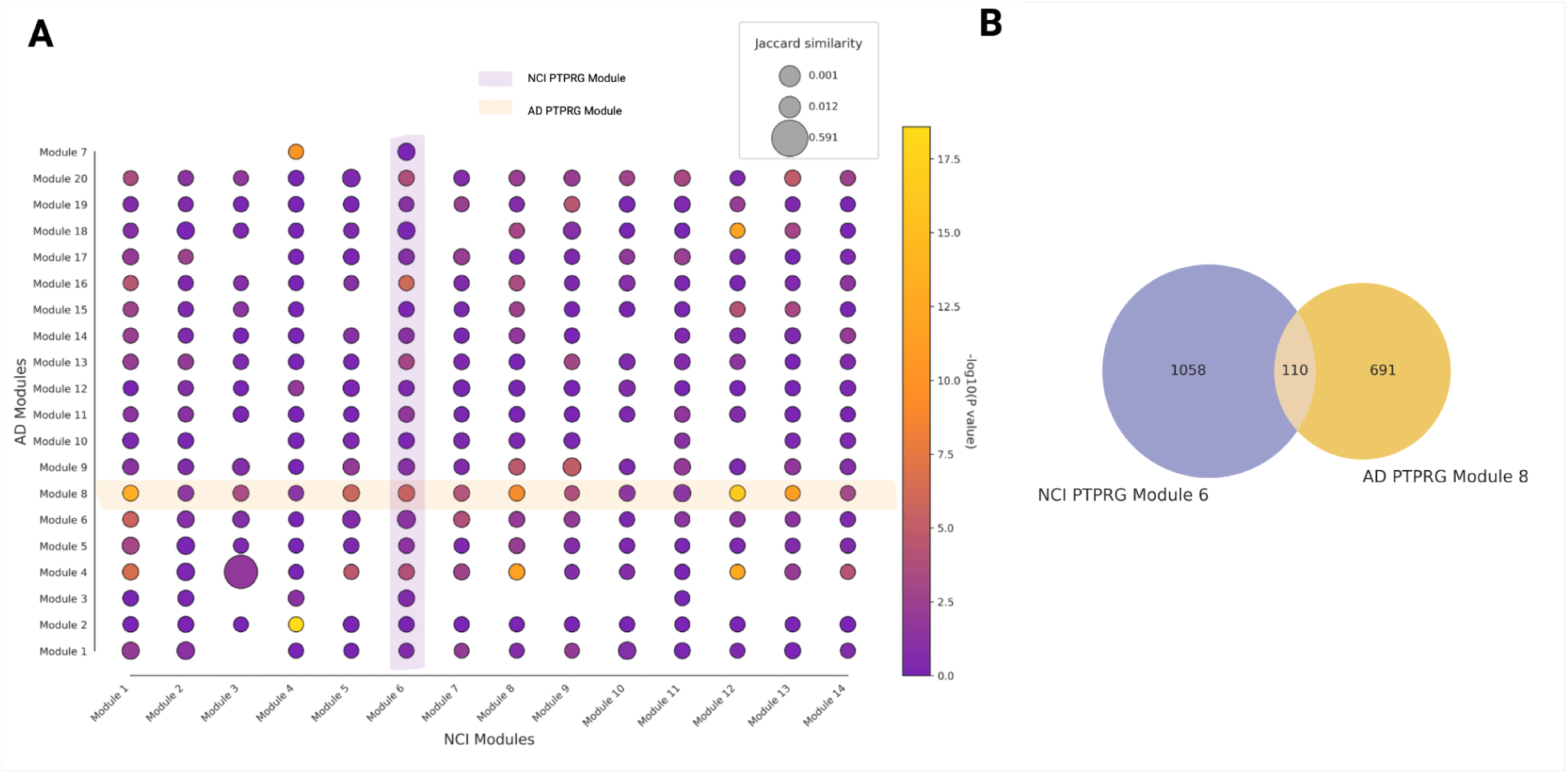
Differential microglial gene co-expression modules in AD and NCI. **A)** Bubble Plot showing distinct modules identified in NCI and AD Microglia. Bubble size reflects the Jaccard similarity (or overlap strength) among the modules, and bubble color encodes the significance of that overlap, with larger, warmer-colored bubbles indicating stronger, more significant overlap. **B)** Venn diagram illustrating overlap and condition-specific genes for PTPRG modules in NCI versus AD microglia

PTPRG was assigned to different modules in each condition-specific network, as illustrated in Figure 5A. In the AD network, its module comprised 801 genes, compared to 1,168 genes in the NCI network. Only 110 genes were shared between the two modules, as shown in the Venn diagram in Figure 5B, indicating that the vast majority of PTPRG co-expression partners differ across conditions and pointing to a substantial reorganization of the gene network surrounding PTPRG in AD. The full gene memberships of the PTPRG-associated modules are provided in Supplementary Table S2.

To characterize the functional roles of genes co-expressed with PTPRG in each condition, we performed GSEA on the AD and NCI modules separately. The two conditions showed different enrichment profiles, reflecting fundamentally distinct microglial states. The AD module yielded 89 significantly enriched pathways dominated by inflammatory and immune signaling, including Toll-like receptor cascades and multiple interleukin pathways, collectively indicative of broad innate immune activation (Figure S3).

In contrast, the NCI module showed a narrower enrichment profile (22 significantly enriched pathways), centered on cellular homeostatic processes such as mTORC1 signaling, oxidative phosphorylation, and protein metabolism (Figure S4). Although some immune-related pathways were also enriched in NCI, these were fewer in number and consistent with homeostatic immune surveillance rather than overt inflammatory activation.

Taken together, these results suggest that PTPRG operates within a fundamentally different gene network context in the two conditions, with AD microglia characterized by widespread inflammatory and immune signaling dysregulation, and NCI microglia embedded in a homeostasis-oriented environment.

The full lists of enriched pathways for the AD and NCI gene sets are provided in Supplementary Table S3.

### Neuronal ligands regulating microglial PTPRG

To identify upstream neuronal signals that could regulate PTPRG expression in microglia, we performed ligand-target analysis using NicheNet (Figure 6). Excitatory and inhibitory neurons shared a largely overlapping set of ligands predicted to regulate microglial PTPRG. Of the top 20 ligands identified for each subtype, 15 were identical, including several AD-associated genes such as APOE and GRN.

**Figure 6.**
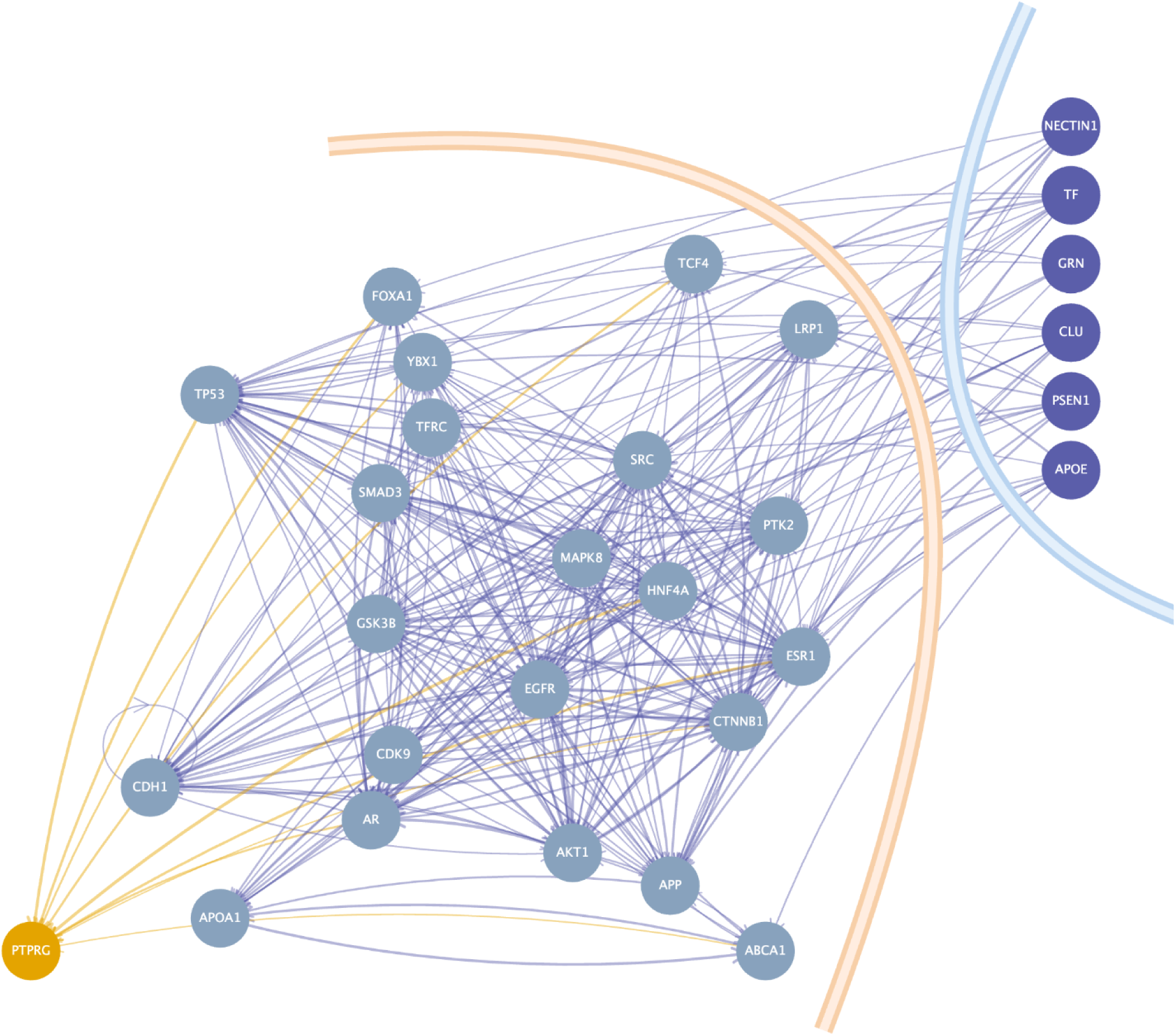
Neuronal-to-microglial ligand-target network. Network graph showing the connections between AD-associated ligands in neurons (blue) and PTPRG in microglia (yellow) through gene regulatory networks. The Network encompasses both excitatory and inhibitory connections.

Each subtype also contributed a small number of distinct ligands: excitatory neurons uniquely included the AD-related gene LPL, while inhibitory neurons uniquely included two major AD-associated genes, PSEN1 and CLU. A complete list of predicted ligand-target interactions and their regulatory weights for both subtypes is provided in Supplementary Table S4.

Consistent with the largely overlapping ligand profiles described above, GSEA performed separately on the top 20 ligands for each neuronal subtype yielded nearly identical enriched pathways, with both subtypes implicating neurodegenerative processes.

### Excitatory and Inhibitory Neurons Engage PTPRG Through Distinct Mechanisms

NicheNet’s reliance on prior knowledge introduces inherent bias into cell-cell communication analysis, and since PTPRG functions as a receptor rather than a ligand, it cannot capture how microglial PTPRG activity may influence neuronal gene expression. We therefore complemented the NicheNet analysis with correlation analysis between microglial PTPRG expression and neuronal gene expression, separately in excitatory and inhibitory neurons. Complete correlation statistics for both excitatory and inhibitory neuron analyses are provided in Supplementary Table S5.

Excitatory neurons showed strong correlation with microglial PTPRG through genes involved in immune activation and antigen presentation (Table 1). GSEA of these genes yielded 20 significantly enriched pathways, predominantly related to immune activation and inflammatory signaling (Figure S5).

**Table 1.** β denotes the regression coefficient reflecting the magnitude and direction of the association with microglial PTPRG expression after accounting for covariates.

| | Gene | Regression coefficient ($\beta$) | Adjusted p-value |
| --- | --- | --- | --- |
| **Excitatory neurons** | RASGEF1B | 2.31 | 0.03 |
|  | INO80D | 2.30 | 0.03 |
|  | SLC26A3 | 2.25 | 0.02 |
|  | DHFR | 1.54 | 0.04 |
|  | SLC27A6 | 1.20 | 0.003 |
| **Inhibitory neurons** | GFAP | 0.147 | 0.006 |
|  | GPIHBP1 | 0.011 | 0.013 |

In contrast, inhibitory neurons showed markedly weaker associations with microglial PTPRG, with only two genes reaching significance. The regression coefficients were approximately 15-fold smaller than those observed for excitatory neurons (maximum β ∼0.15 vs. ∼2.3), indicating weaker transcriptional coordination between PTPRG-expressing microglia and inhibitory neurons (Table 1). The pathway enrichment profile was also qualitatively distinct from that of excitatory neurons, with 14 significantly enriched pathways centered on lipid metabolism, cellular stress response and protein homeostasis (Figure S6).

Collectively, these results indicate that excitatory and inhibitory neurons are coupled to microglial PTPRG activity through fundamentally different biological processes, suggesting that the two neuronal subtypes may contribute to AD-associated microglial dysregulation through distinct mechanisms.

## Discussion

This study demonstrates that glial transcriptional signatures carry the strongest discriminative signal between AD and non-cognitively impaired individuals in human brain snRNA-seq data, and identifies microglial PTPRG as a central molecular hub integrating neuronal signals and inflammatory dysregulation in disease pathology. By systematically evaluating all 63 possible cell-type combinations through a pseudobulk classification framework, we provide the first unbiased, data-driven ranking of cell-type contributions to AD-associated transcriptional changes, with microglia and astrocytes together yielding the best performing model. The resulting signatures generalize robustly to an independent cohort, supporting their biological reproducibility across datasets with different brain region sampling, library chemistry, and demographic composition.

The primacy of glial cell types in AD classification is consistent with a growing body of evidence implicating microglia and astrocytes as active drivers rather than passive responders in disease pathophysiology. Large-scale snRNA-seq studies have identified disease-associated microglial and astrocytic subpopulations that emerge in response to amyloid and tau pathology and contribute to neuroinflammation, synaptic dysfunction, and neurodegeneration (Mathys et al., 2019; Habib et al., 2020; Olah et al., 2020; Green et al., 2024). Our results extend these findings by demonstrating that the transcriptional divergence between AD and NCI is most concentrated in these two cell types when evaluated through a supervised classification lens, and that this signal is sufficiently robust to generalize across cohorts. While prior computational approaches either analyzed cell types independently, potentially missing coordinated cross-cell-type signals (He et al., 2024), or pooled all cells without preserving cell-type identity (Verlaan et al., 2025), our pseudobulk-per-cell-type strategy retains cell-type specificity while operating at the sample level, avoiding the cell-level label assignment problem that complicates direct application of supervised learning to single-cell data. Imaging-based machine learning has previously discriminated AD from control at the single-cell level (Muñoz-Castro et al., 2022), but to our knowledge no prior study has formally quantified the relative discriminative contribution of each major brain cell type through supervised classification with systematic combinatorial evaluation of all possible combinations.

Among the 72 genes selected by the final model, microglial PTPRG emerged as the strongest individual predictor of AD. PTPRG encodes a receptor-type protein tyrosine phosphatase that regulates cell signaling by dephosphorylating tyrosine residues on intracellular substrates, and receptor-type phosphatases of this class are established negative regulators of innate immune activation, modulating inflammatory signaling thresholds by opposing kinase-driven cascades. The strong downregulation of PTPRG in AD microglia, combined with WGCNA evidence that its co-expression network shifts from a homeostatic, metabolically oriented context in NCI to a broadly inflammatory one in AD, suggests that loss of PTPRG-mediated phosphatase activity may contribute to the failure of inflammatory resolution in disease-associated microglia. This interpretation is supported by the GSEA findings showing enrichment of Toll-like receptor cascades and multiple interleukin signaling pathways in the AD PTPRG module, in contrast to the narrower homeostatic enrichment profile centered on mTORC1 signaling, oxidative phosphorylation, and protein metabolism observed in NCI. The fact that only 110 of the genes co-expressed with PTPRG are shared between the AD and NCI networks, out of modules comprising 801 and 1168 genes respectively, indicates a near-complete reorganization of the regulatory context in which PTPRG operates, rather than a simple change in expression level.

The NicheNet analysis positions PTPRG as a convergence point for upstream neuronal signals, with established AD risk genes including APOE, GRN, PSEN1, and CLU among the top predicted ligands from both excitatory and inhibitory neurons. This pattern suggests that disease-relevant neuronal dysfunction feeds into microglial PTPRG regulation through ligand-receptor mechanisms, linking neuronal AD pathology to microglial inflammatory state through a shared molecular node. The differential coupling of the two neuronal subtypes to microglial PTPRG adds further specificity to this model. Excitatory neurons showed strong transcriptional coordination with PTPRG through immune activation and antigen presentation pathways, with regression coefficients approximately 15-fold larger than those observed for inhibitory neurons, which showed weaker coupling focused on lipid metabolism and cellular stress response. This asymmetry is consistent with the selective vulnerability of excitatory circuits in AD and with previous observations that excitatory neuron subtypes are preferentially depleted in AD-affected brain regions (Leng et al., 2021; Mathys et al., 2023), suggesting that excitatory neurodegeneration and microglial inflammatory dysregulation are transcriptionally coupled through PTPRG, while inhibitory neurons engage microglia through metabolic rather than immune channels.

The astrocytic component of the model complements this picture. The top astrocytic features, including IGFBP5 and SORCS3, and the enrichment of HSF1-mediated chaperone pathways in the astrocyte gene set point to a coordinated proteostasis stress response in AD astrocytes, consistent with the established role of astrocytes in protein quality control and their reactive transformation under sustained neuroinflammation. The two glial programs likely interact: chronic microglial inflammatory signaling plausibly contributes to the proteostatic burden that astrocytes attempt to manage, linking the two cell-type-specific signatures into a coherent model of coordinated glial dysfunction in AD.

The graded distribution of predicted AD probabilities for MCI+P samples, falling between those of extreme AD and NCI cases, indicates that the classifier captures a biologically meaningful disease gradient rather than a binary distinction. This is consistent with the conceptualization of AD as a continuum of accumulating pathology (Aisen et al., 2017) and suggests that glial transcriptional signatures track disease severity progressively, offering potential utility for staging patients along the disease spectrum rather than simply separating diagnostic categories.

Several experimental directions follow directly from these findings. Tri-culture systems comprising neurons, astrocytes, and microglia have been established to recapitulate key AD pathological features including microglia-driven neuroinflammatory responses mediated through astrocyte-microglia crosstalk (Park et al., 2018; Luchena et al., 2022). Within this framework, CRISPR-mediated perturbation of PTPRG specifically in microglia would enable mechanistic dissection of its cell-autonomous and non-cell-autonomous effects on neuronal viability and inflammatory signaling, directly testing the model proposed here. Longitudinal single-cell data tracking these signatures from early pathological stages could further determine whether PTPRG network reorganization precedes clinical symptoms and define therapeutic windows for intervention. More broadly, the pseudobulk classification framework developed here is not specific to AD and is directly applicable to other neurodegenerative diseases where cell-type-resolved transcriptional data are available, offering a generalizable strategy for identifying the cellular drivers of complex brain pathology.

Beyond its immediate mechanistic implications, this work has broader relevance for the field. The demonstration that a small set of glial transcriptional features, derived from routinely available snRNA-seq data and validated across independent cohorts, can reliably separate AD from cognitively normal individuals opens a path toward transcriptomics-informed patient stratification. As single-cell atlases of the human brain continue to expand in scale and diversity, frameworks of the kind developed here could support the identification of molecularly defined patient subgroups with distinct glial activation profiles, potentially informing trial enrichment strategies or the prioritization of cell-type-specific therapeutic targets. The convergence of the classification signal on PTPRG and a small number of co-regulated immune and proteostatic genes further suggests that AD-associated glial dysfunction may be more tractable than the complexity of the disease has historically implied, with a limited set of molecular nodes driving much of the transcriptional divergence between disease and health.

This study has a number of methodological strengths. The systematic evaluation of all 63 cell-type combinations provides an unbiased assessment of cellular contributions to disease classification that does not assume any prior biological knowledge about which cell types are most relevant. The use of consistent feature selection across cross-validation folds ensures that the final gene set is stable and reproducible rather than the product of a single fit. Cross-cohort validation on an independent dataset with different brain region sampling, library chemistry, and demographic composition provides strong evidence that the identified signatures reflect genuine biology rather than dataset-specific technical artefacts. The integration of four complementary analytical approaches, supervised classification, WGCNA, NicheNet, and correlation analysis, converges on a coherent and interpretable biological model centered on PTPRG, strengthening confidence in this finding beyond what any single method could support. Finally, the graded prediction of intermediate disease states without retraining suggests that the model captures continuous molecular variation along the disease spectrum, extending its potential utility beyond binary classification.

Several limitations should be considered alongside these strengths. Pseudobulk aggregation reduces the sparsity inherent in single-cell data but can conflate gene expression shifts with changes in cell-type composition within a sample, since both produce differences in aggregated profiles. Training on extreme phenotypes enhances class separation but may reduce sensitivity to the more subtle transcriptional changes present in early-stage disease. Cohort-specific technical factors including brain region sampling, library chemistry, and demographic composition can influence signals despite cross-cohort alignment steps, and L1 regularization coefficients provide a useful ranking of feature importance but their absolute magnitudes depend on preprocessing and scaling choices and should not be interpreted as effect sizes. The SEAD validation cohort exhibited substantial class imbalance (39 AD vs. 9 NCI), which limits the precision of performance estimates in that dataset, particularly for sensitivity, and replication in a more balanced independent cohort would strengthen confidence in these findings. Finally, all analyses are based on postmortem brain tissue, which captures a late disease state and precludes inference about the temporal ordering of the molecular events identified here.

Taken together, this work provides a systematic, multi-layered characterization of the cellular and molecular architecture of AD transcriptional pathology. By establishing glial signatures as the dominant discriminative signal, identifying PTPRG as a convergence node linking neuronal AD risk biology to microglial inflammatory state, and demonstrating cross-cohort generalizability, these findings advance both the mechanistic understanding of AD and the methodological toolkit for interrogating complex neurodegenerative diseases at single-cell resolution. PTPRG represents a compelling candidate for experimental follow-up and, given its role as a negative regulator of inflammatory signaling, a potential target for therapeutic strategies aimed at restoring microglial homeostasis in AD.

## Acknowledgements

This study was supported by a starting grant from the Chalmers Area of Advance Health Engineering to AP. The data handling was enabled by resources provided by the National Academic Infrastructure for Supercomputing in Sweden (NAISS), partially funded by the Swedish Research Council through grant agreement no. 2022-06725.

The results published here are in whole or in part based on data obtained from the AD Knowledge Portal (https://adknowledgeportal.org/). Study data were generated from postmortem brain tissue provided by the Religious Orders Study and Rush Memory and Aging Project (ROSMAP) cohort at Rush Alzheimer’s Disease Center, Rush University Medical Center, Chicago. This work was supported in part by the Cure Alzheimer’s Fund, NIH grants AG058002, AG062377, NS110453, NS115064, AG062335, AG074003, NS127187, MH119509, HG008155 (M.K.), RF1AG062377, RF1 AG054321, RO1 AG054012 (L.-H.T.) and the NIH training grant GM087237 (to C.A.B.). ROSMAP is supported by P30AG10161, P30AG72975, R01AG15819, R01AG17917. U01AG46152, U01AG61356.

## Data Availability

The datasets described in this study are available at the AD Knowledge Portal (https://adknowledgeportal.org), under the following accession IDs: syn52293433, syn26223298.

**Code Availability** https://github.com/Polster-lab/celltype-resolved-ad Author contributions

**Conceptualization:** D.A., J.G., A.P. **Data curation:** D.A., A.M., N.P.M. **Formal analysis:** D.A., A.M., N.P.M. **Funding acquisition:** A.P. **Investigation:** D.A., A.M., N.P.M. **Methodology:** D.A., A.P. **Project administration:** A.P. **Software:** D.A., A.M. **Supervision:** J.G., A.P. **Validation:** D.A., A.M. **Visualization:** D.A., A.M., N.P.M. **Writing:** D.A., A.M., N.P.M., E.K., J.G., A.P.

## Supplementary Figures

**Figure S1.**
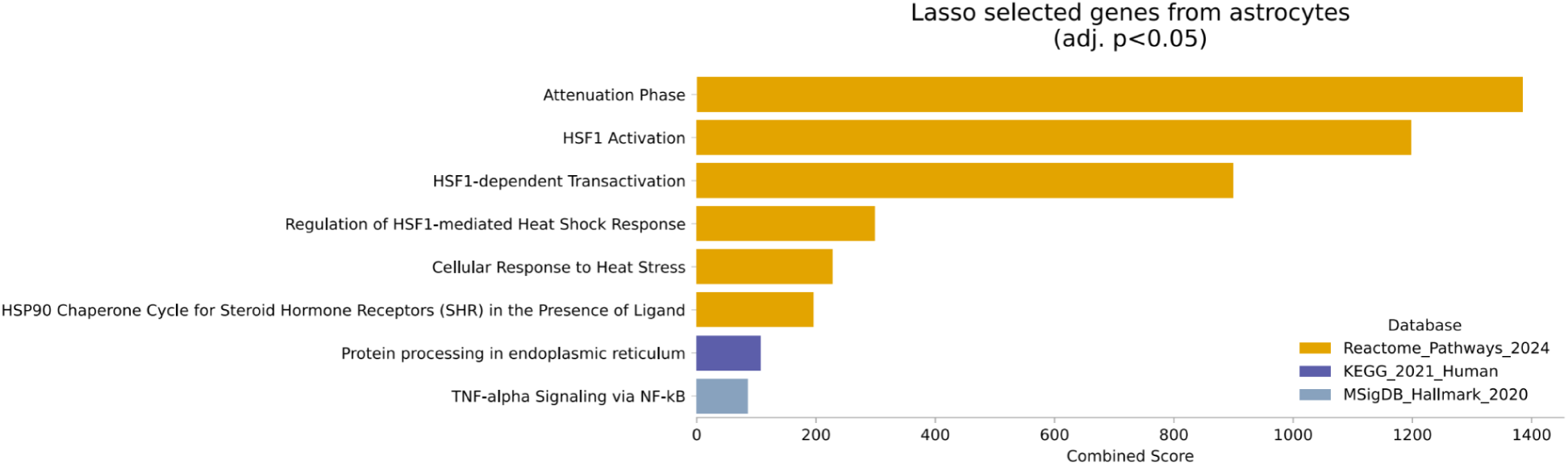
Enriched Biological Pathways of LASSO-Selected Astrocyte Genes (Adjusted p < 0.05)

**Figure S2.**
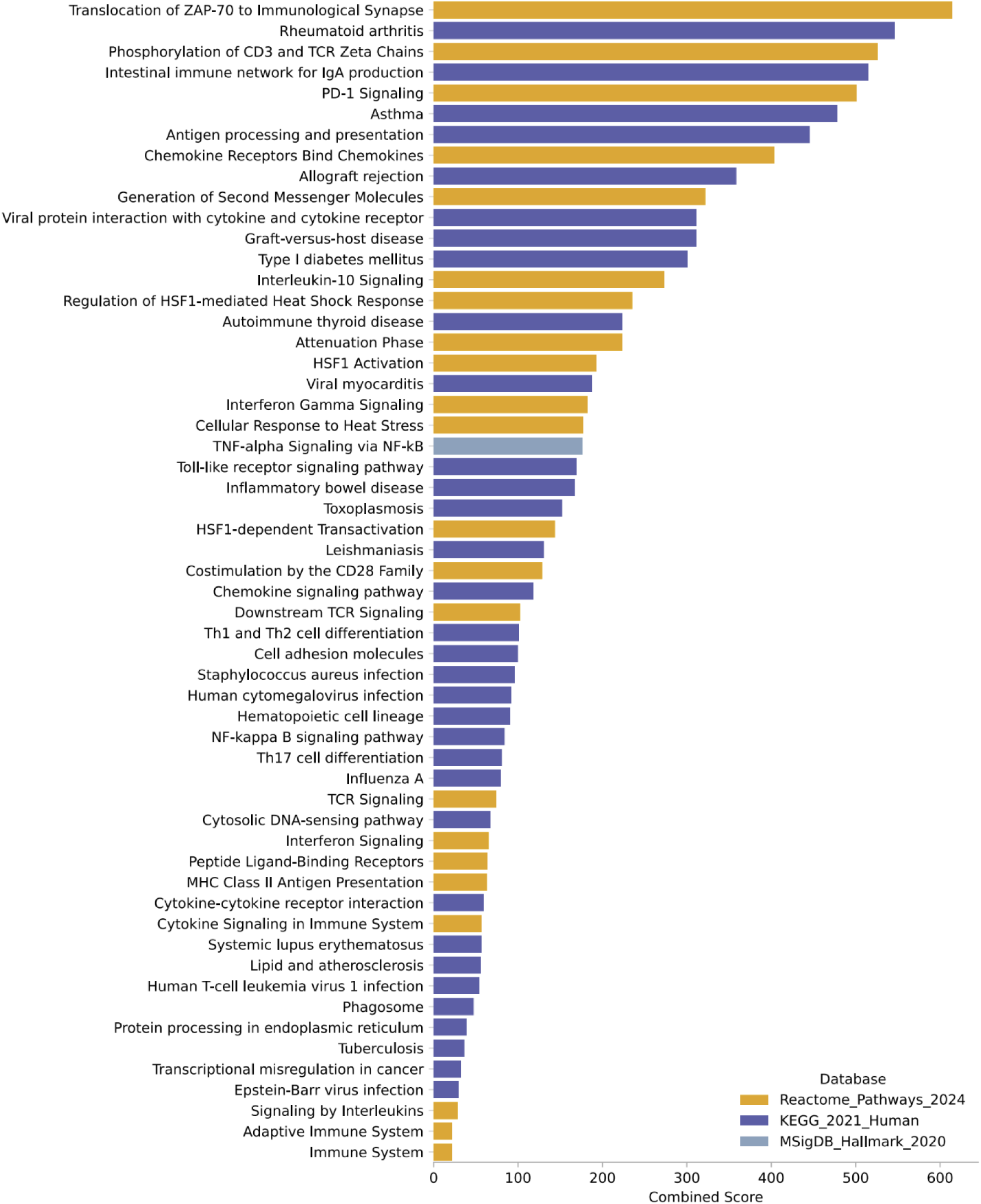
Enriched Biological Pathways of LASSO-Selected Microglia Genes (Adjusted p < 0.05)

**Figure S3.**
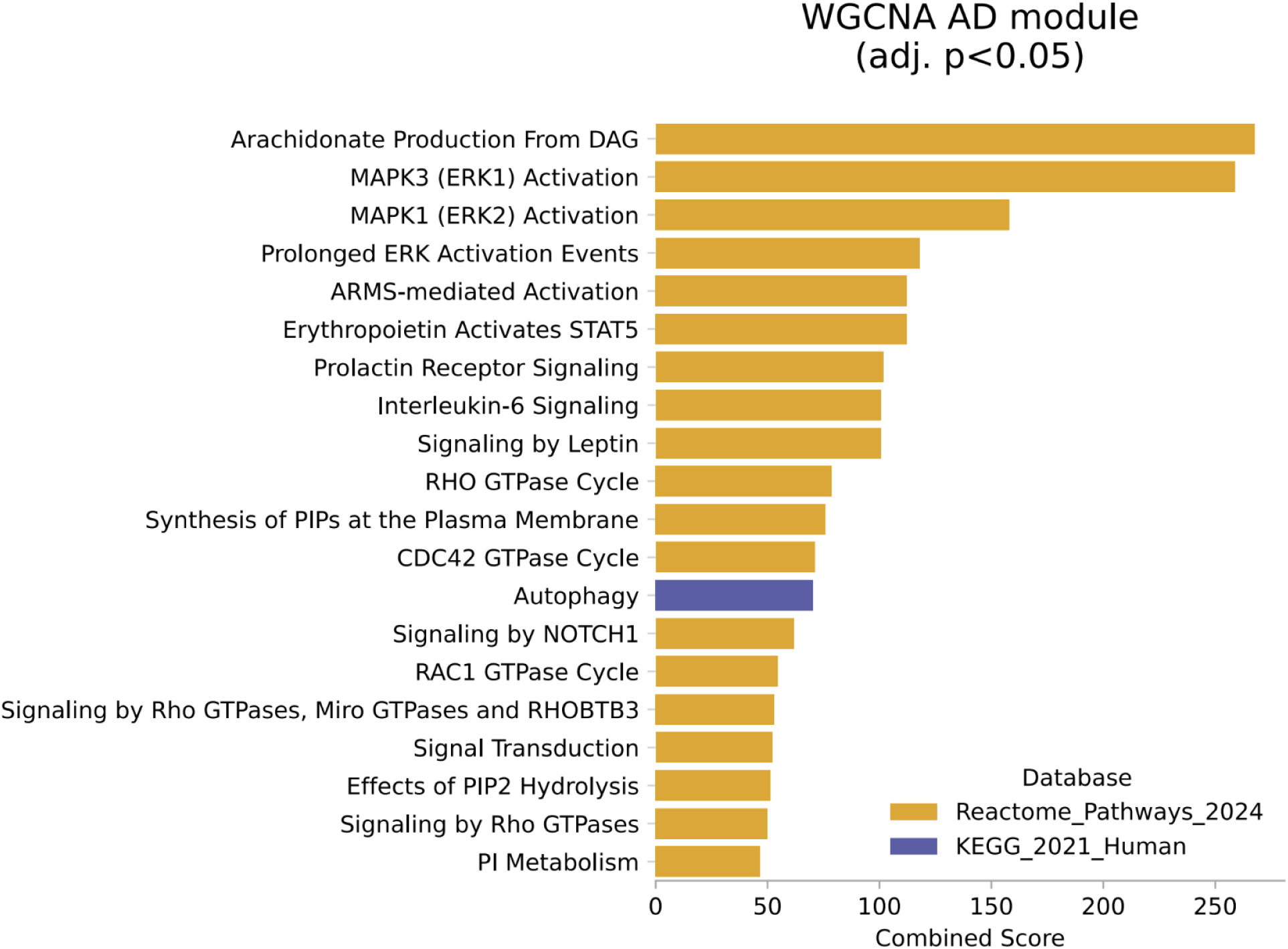
Enriched Biological Pathways of WGCNA AD module Genes (Adjusted p < 0.05)

**Figure S4.**
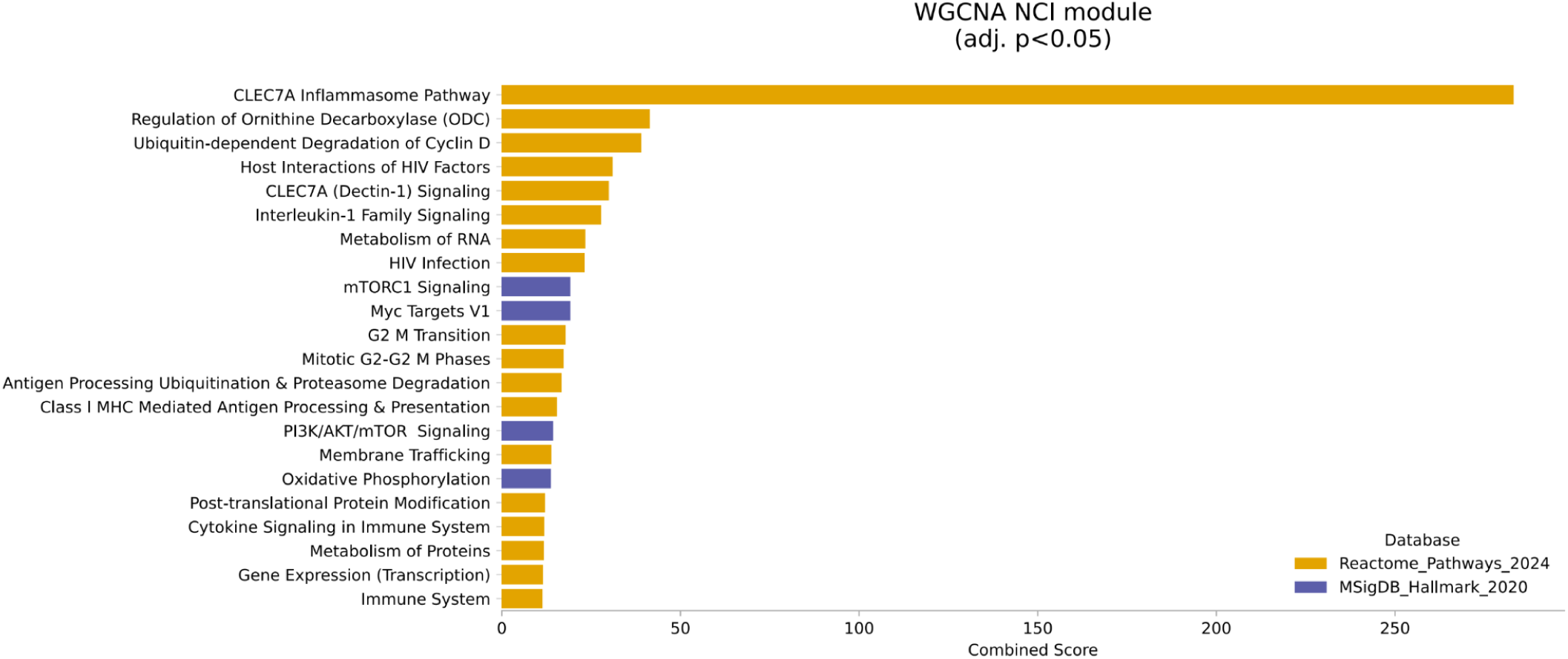
Enriched Biological Pathways of WGCNA NCI module Genes (Adjusted p < 0.05)

**Figure S5.**
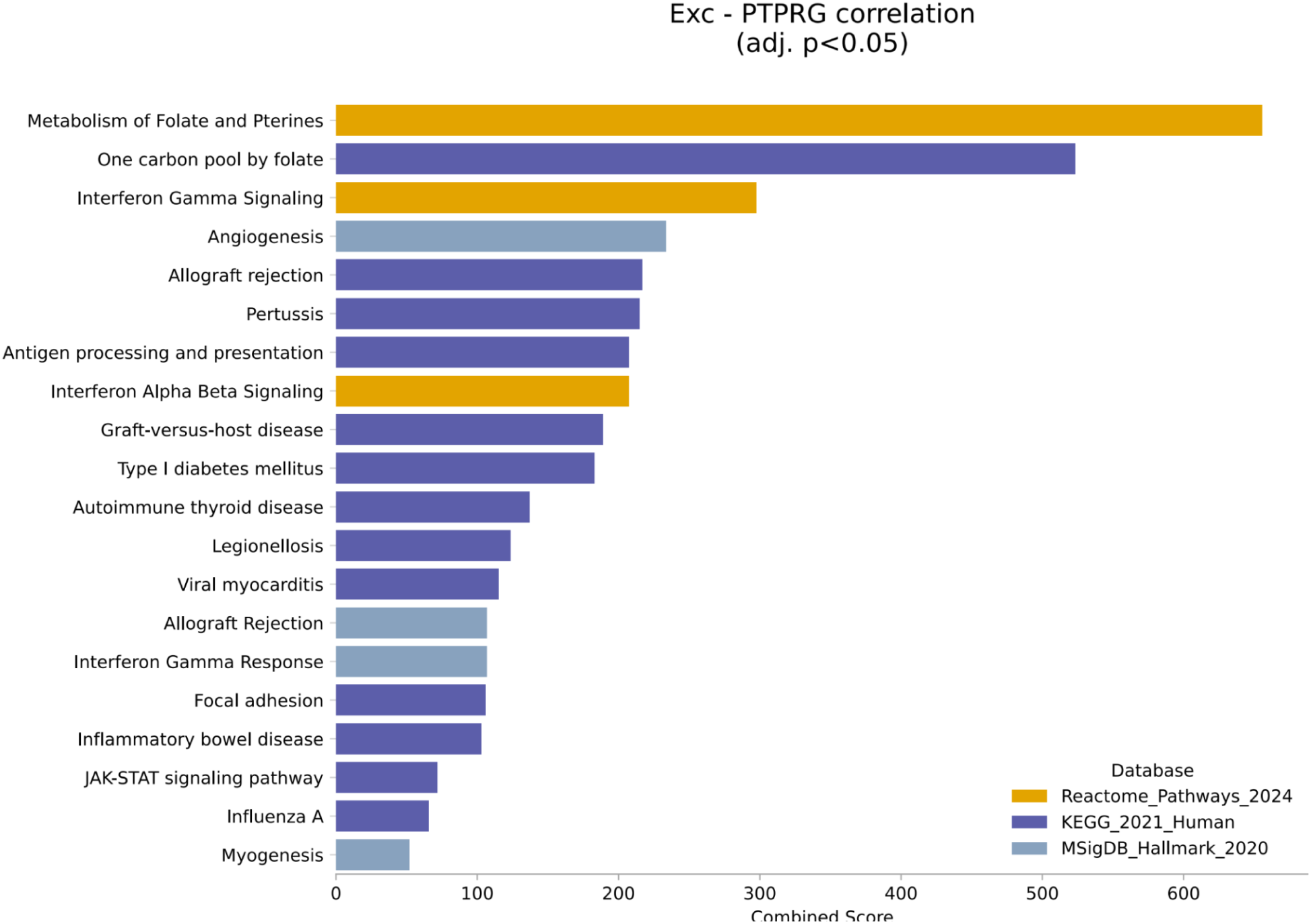
Pathway Enrichment of Excitatory Neuron Genes Correlated with Microglial PTPRG (Adjusted p < 0.05)

**Figure S6.**
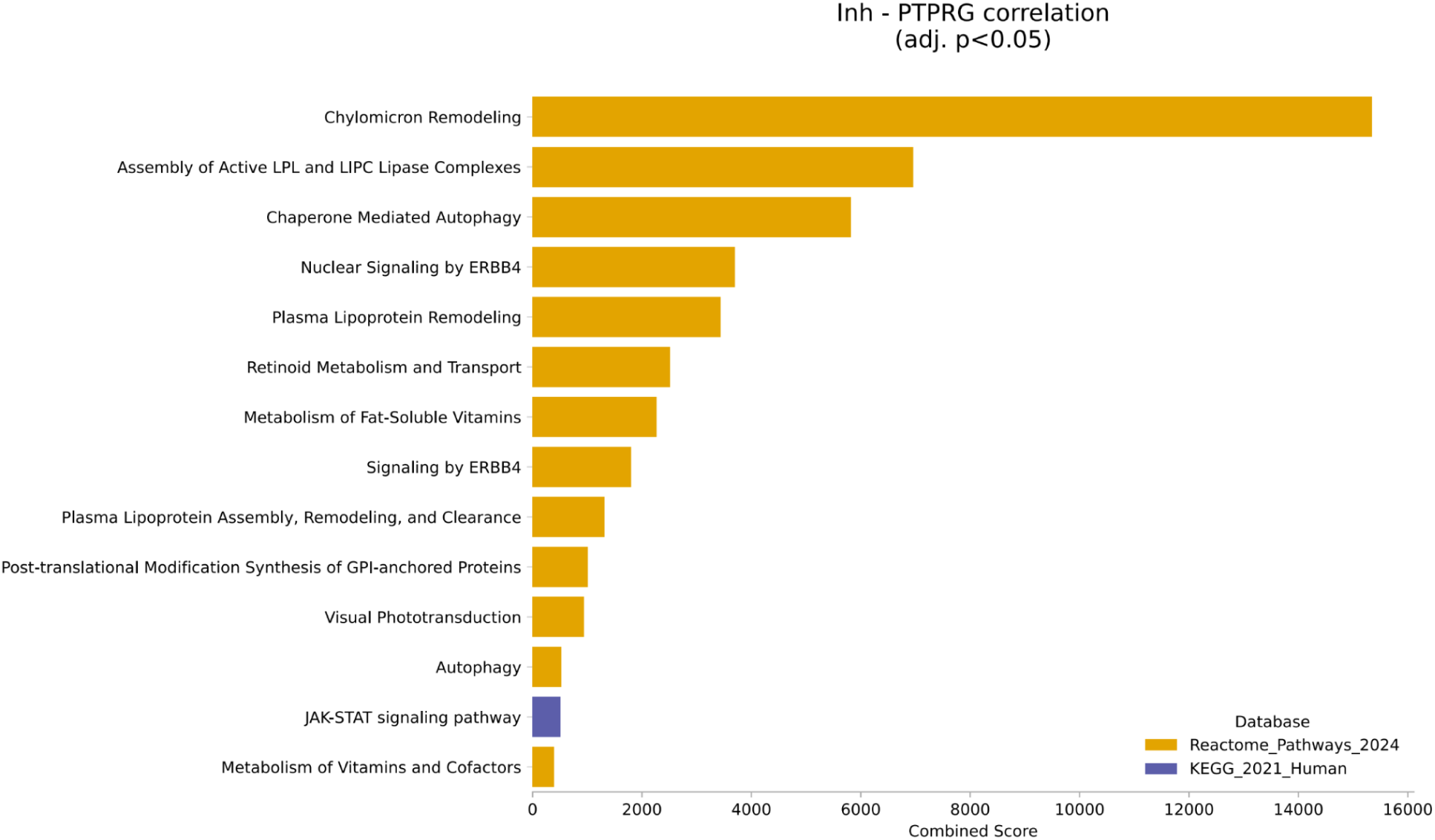
Pathway Enrichment of Inhibitory Neuron Genes Correlated with Microglial PTPRG (Adjusted p < 0.05)

## Supplementary Tables

**Table S1. Optimal cell-type combination for AD classification**

**Table S2. Full gene memberships of the PTPRG-associated modules in AD and NCI**

**Table S3. Enriched pathways for the genes in AD and NCI PTPRG modules.**

**Table S4. Predicted Neuronal Ligand–Target Interactions Regulating Microglial PTPRG Identified by NicheNet**

**Table S5. Correlation statistics for both excitatory and inhibitory neuron analyses**

